# Germline Selection by Meiosis Defends the Transmission of defective Mitochondria with mtDNA variants

**DOI:** 10.1101/2020.11.10.377390

**Authors:** Hongying Sha, Yimin Yang, Sanbao Shi, Dongmei Ji, Jianxin Pan

**Author notes:** Co-first author (Hongying Sha, Yimin Yang, Sanbao Shi).

## Abstract

Germline selection of mtDNA is vital in maternal inheritance of mtDNA, as it can eliminate severe mtDNA mutations. However, current evidence concerning germline selection at meiosis level comes from incomplete mtDNA sequencing in human first polar body (PB1), which lacks persuasion. Here, we found various variants, including pathogenic mutation sites, present on whole genome of mtDNA in human PB1 compared with its oocyte. And that PB1 mitochondria with mtDNA variants were defective. Afterwards, to further explore how mitochondria enter PB, the defective mitochondria transfer in mouse germline, including cumulus-oocyte-complexes at germinal vesicle and matured oocytes stage. It confirmed that in the first and second meiosis, active purification selected defective mitochondria into PB1 and PB2. Thus, twice meiosis is the last defense system for purifying selection of mtDNA mutations during oogenesis, which also demonstrated that PB1 and PB2 would be final destination of deleterious mtDNA mutations in germline selection.

## INTRODUCTION

Mitochondria and mitochondrial DNA (mtDNA) are inherited exclusively through the maternal lineage of mammals (Babayev et al., 2015). Thus an oocyte containing mutant mtDNA is likely to give birth to affected offspring (Brown et al., 2006). According to statistics, the incidence of mtDNA disease is high, at least 1 in 5000 (Schaefer et al., 2004, 2008). Although mitochondrial DNA has numerous mutation sites, the mtDNA mutation-related diseases we found come from mutations at that few sites. In other words, only a few mtDNA mutations account for the majority of mtDNA disease. This situation suggests that the severe mutations can be selectively eliminated in the female germ line (Lieber et al., 2019, Wei et al., 2019, Fan et al., 2008, Stewart et al., 2008, Shoubridge et al., 2008). The germline selection may contain four levels: genome level (mitochondrial genetic bottleneck), organelle level (autophagocytosis), cellular level (apoptosis in preovulatory follicles) (Stewart et al., 2008, Fan et al., 2008), and meiosis level (Fanti et al., 2017, Gianoarli et al., 2014).

However, in the study to date, selection by meiosis still lacks direct experimental support Existing evidence support for germline selection by meiosis only comes from incomplete sequencing of mtDNA genome in human first polar body (PB1) (De Fanti et al., 2017, Gianoarli et al., 2014). Moreover, since mtDNA itself is polymorphic, the variants cannot be convincing evidence of mtDNA selection. Furthermore, sequencing is not the most intuitive evidence for meiosis selection of mtDNA variants. As we know, meiosis is a visual and traceable biological event under a microscope. In light of these characteristics, we can observe mtDNA mutations selected during meiosis by labeling mitochondria, such as live cell fluorescent probes, under a microscope.

In the present study, to address these issues, we compared variants of the entire mtDNA genome in PB1 and its oocytes. Then, the events of mtDNA selection by meiosis were tracked via defective mitochondrial transfer in mouse germline. Our study provides intuitive evidence to support the existence of meiosis selection against mtDNA variants.

## RESULTS

### Accumulation of mtDNA variants present in human PB1 relative to its oocyte

The donors were twelve women with ages ranging from 25 to 31, named W_1_~W_12_. A total of 18 matured oocytes were obtained from 25 metaphase I oocytes according the rotuine procedure in our lab (Zou et al. 2019). Correspondingly, oocytes denoted by these women were also named in the order of O_1_ to O_12_. Among them, W_1_ to We donated one oocyte, respectively. Oocytes denoted by W_1_~W_6_ were used as mtDNA sequencing of single cell, including single PB1 and oocyte. Whole Genome Amplification (WGA) was applied for amplifying a single PB1 and oocyte genome (Figure. 1A). Meanwhile, in order to verify whether WGA alters the genome sequence, two mixed samples (12 polar bodies and 12 oocytes donated by W_7_~W_12_ women) were directly lysed to obtain genomes without WGA. Human mitochondrial genomes of all samples were then amplified in 9, 16 or 23 overlapping PCR fragments (Table. S1, Figure. 1B). Subsequently, Sanger sequencing was manipulated on a total of 14 mitochondrial genomes. The sequencing coverage of all single samples and mixed samples (97.39% ~ 99.92%). Percentage mapped to the whole mtDNA genome was significantly higher than other reported studies, which had sequencing coverage of 9-95% and 69.7%, respectively (Fanti et al., 2017, Gianoarli et al., 2014). The ranges of total variants were 9 to 51 (Fig. 1C). Various variants present through the whole mtDNA genomes in PB1 compared to its oocytes (Fig. 1C, Table.S2, Spreadsheat S1).

**Figure. 1.**
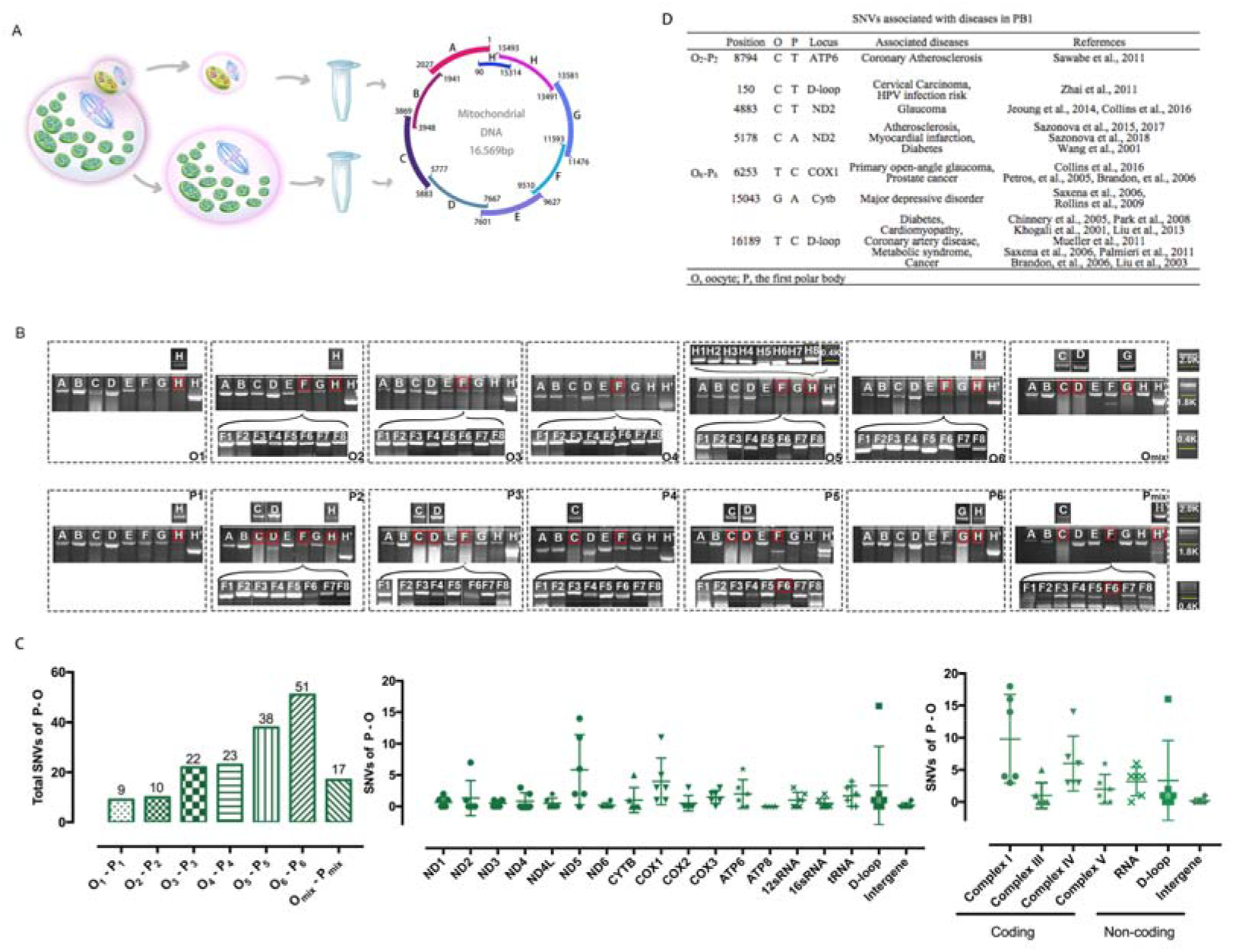
Accumulation of mtDNA variants present in human PB1 relative to its oocyte. A. Schematic charts of experimental design of sequencing mtDNA variants in human PB1 and oocyte. B. Amplicons (A-H’) and (Fl-F8, Hl-H8) of mitogenomes from each single oocyte and PB1. An electrophoresis map represents each amplified fragment of a single oocyte and PB1 sample. Red boxes marked those fragments that failed to be amplified or sequenced. M, size marker. C. Comparison of total mtDNA variants between PB1 and its sister oocyte. Distribution of mtDNA variants in different regions compared to PB1 and its sister oocyte. Total mtDNA variants in coding and non-coding regions compared between PB1 and its sister oocyte. D. SNVs associated with diseases in mtDNA.

Interestingly, in two PB1s from O_2_ and O_6_, we found six and one pathogenic mutation sites respectively associated with Cervical Cancer, HPV infection risk (Zhai et al., 2011), Glaucoma (Jeoung et al., 2014, Collins et al., 2016), atherosclerosis (Sazonova et al., 2015, 2017), myocardial infarction (Sazonova et al., 2018), diabetes (Wang et al., 2001, Chinnery et al., 2005, Park et al., 2008), primary open-angle glaucoma (Collins et al., 2016), Prostate Cancer (Petros, et al., 2005, Brandon, et al., 2006), Major Depressive Disorder (Saxena et al., 2006, Rollins et al.,2009), Cardiomyopathy (Khogali et al., 2001, Liu et al., 2013), coronary artery disease (Mueller et al., 2011), Metabolic Syndrome (Saxena et al., 2006, Palmieri et al., 2011), cancer (Brandon, et al., 2006, Liu et al., 2003), Coronary Atherosclerosis (Sawabe et al., 2011) (Fig. 1D). These pathogenic mutation sites strongly suggested that mtDNA mutations would be accumulated within PB1. Thus, we then compared the mitochondrial characteristics of human PB1 and its sister oocyte.

### Human PB1 Contained defective Mitochondria than its Sister Oocyte

It has been demonstrated that TMRE and DilC_1_ (5) accumulate in active mitochondria due to their relative negative charge. As a result, inactive mitochondria have decreased membrane potential and fail to sequester TMRE and DilC_1_ (5) (Chazotte 2011). And Rhod2 and Fluo-3 are the most commonly used fluorescent stains to detect inactive mitochondria reflected by intracellular calcium concentration (Orrenius 2015). Thus, TMRE and DilC_1_ (5), Rhod2 and Fluo-3 mitochondrial membrane potential assay kit were used to study the mitochondria activation in human PB1s and its oocyte. In experiments, we found that TMRE and DilC_1_ 5 were absent in human PB1 while TMRE and DilC_1_5 were abundant in human oocyte.(Figure. 2A). The fluorescence intensity for TMRE and DilC_1_5 in human PB1 was significantly lower than that of their sister oocytes (Figure. 2B, P<0.05 (DilC15 p=0.001, TMRE p=0.0263)). However, high levels of Rhod2 and Fluo-3 accumulated in human PB1 while very low level of signals were found in human oocyte (Figure. 2A). The fluorescence intensity for Rhod2 and Fluo-3 in human PB1 was significantly higher than that of their sister oocytes (Figure. 2B, P<0.05 (Rhod2 p=0.0237, Fluo-3: p=0.0117)).

**Figure. 2.**
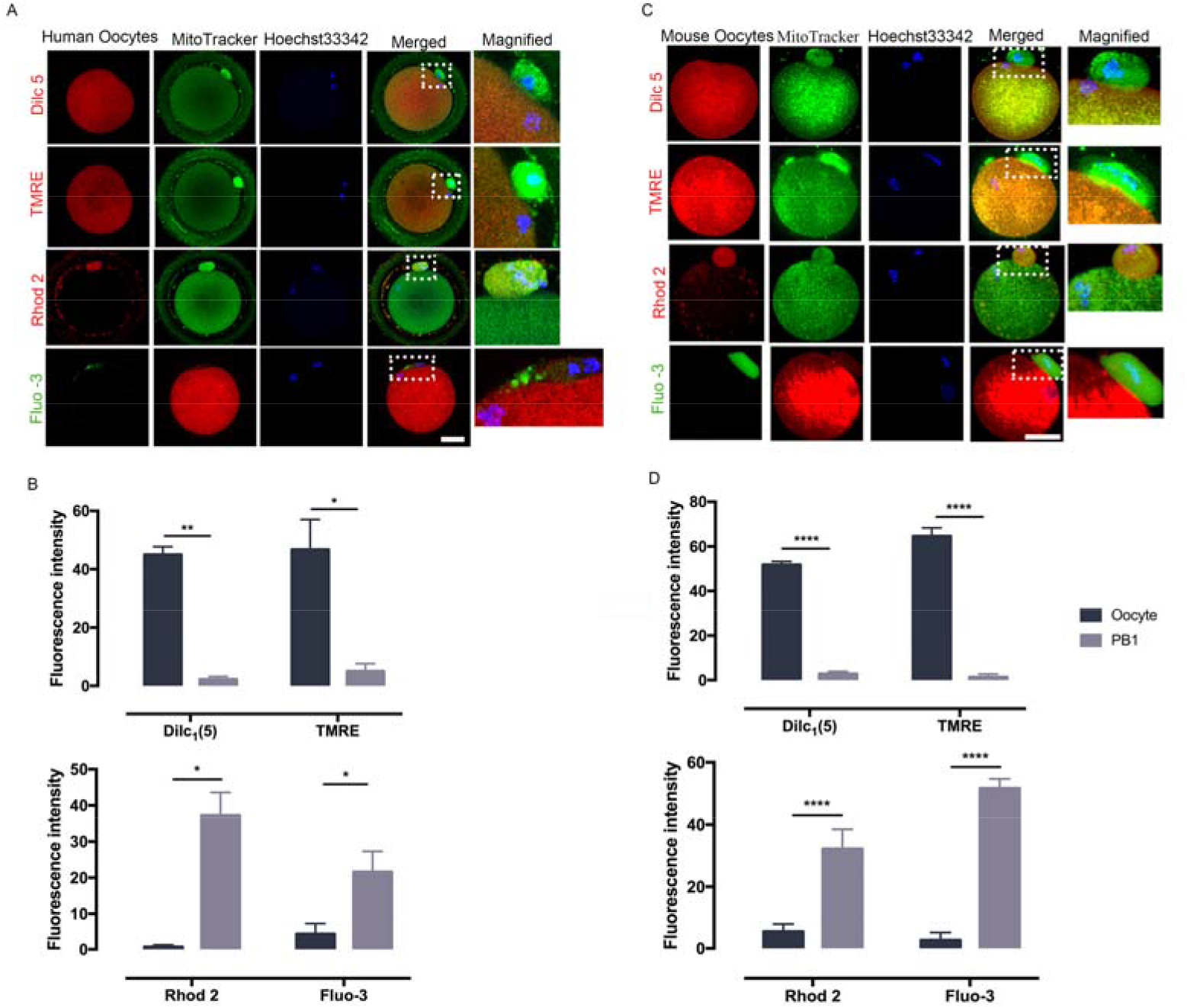
Human and mouse PB1 Contain defective mitochondria than its Sister Oocyte. A. Mitochondrial membrane potential was detected with TMRE and DilC_1_(5), Rhod2 and Fluo-3 in human PB1 and oocytes. B. The relative fluorescence intensity of human PB1 to its sister oocyte was quantified and the difference significance was evaluated using paired t test. C. Mitochondrial membrane potential was detected with TMRE and DilC_1_(5), Rhod2 and Fluo-3 in mouse PB1 and oocytes. D. The relative fluorescence intensity of mouse PB1 to its sister oocyte was quantified and the difference significance was evaluated using paired t test. In TMRE and DilC_1_(5), Rhod2 detection, red color represented TMRE and DilC_1_(5), Rhod2, green color marked mitochondria in PB1 and oocyte. In Fluo-3 detection, green color showed Fluo-3, red color stained mitochondria in PB1 and oocyte, ****p < 0.0001, ***p < 0.001, **p < 0.01, *p < 0.05. Error bars indicate SD, with the mean value, n=3 per group for human, n = 6 per group for mouse. Scale bar=40μm.

These mitochondrial differences between PB1 and its sister oocyte suggested that defective mitochondria with mtDNA mutations was enriched in PB1 compared to its sister oocyte. How was these defective mitochondria with mtDNA mutations enriched in PB1? Could defective mitochondria also be enriched in PB2? To reveal these, we conducted the defective mitochondria transfer in mouse germline, including cumulus-oocyte-complexes at germinal vesicle and oocytes after meiosis I.

### The Defective Mitochondria Transfer in GV COCs Revealed Meiosis I actively Selected Defective Mitochondria into PB1

We first monitored the properties of mitochondria in mouse PB1s and its oocytes using TMRE and DilC_1_ (5), Rhod2 and Fluo-3 mitochondrial membrane potential assay kit. We found that, like human PB1, mitochondria were defective in mouse PB1s (Figure 2C). The fluorescence intensity for TMRE and DilCl5 in mouse PB1 was significantly lower than that of their sister oocytes (Figure.2D, P<0.0001). The fluorescence intensity for Rhod2 and Fluo-3 in mouse PB1 was significantly higher than that of their its sister oocytes (Figure.2D, P<0.0001). Then we transferred defective mitochondria from PB1 into cumulus-oocyte-complexes at germinal vesicle (GVCOCs) or broken GVCOCs to observe the destination of defective mitochondria during the first meiotic maturation (Figure 3A, B, Movie.S1). We presumed the transferred mitochondria would have three destinations: completely retained in the matured oocyte, completely extruded into PB1 of the matured oocyte, or some mitochondria retained in the oocyte and some into PB1. (Figure. 3A). After manipulating and maturing in vitro for 10 hours, 23 oocytes with PB1s were obtained from 60 GVCOCs. We found that 60.87% (14/23) oocytes completely extruded the transferred mitochondria into their PB1s, while the remaining nine oocytes retained the implanted mitochondria in the cytoplasm (Figure. 3B, C, Table.S3). For broken GVCOCs group, 15 oocytes with PB1s were obtained from 25 broken GVCOCs after defective mitochondria manipulating and maturing in vitro for 6 hour. However, none of the 15 oocytes extruded the transferred mitochondria into their PB1s. Instead, all transferred mitochondria were retained in the oocytes (Table.S3).

**Figure. 3.**
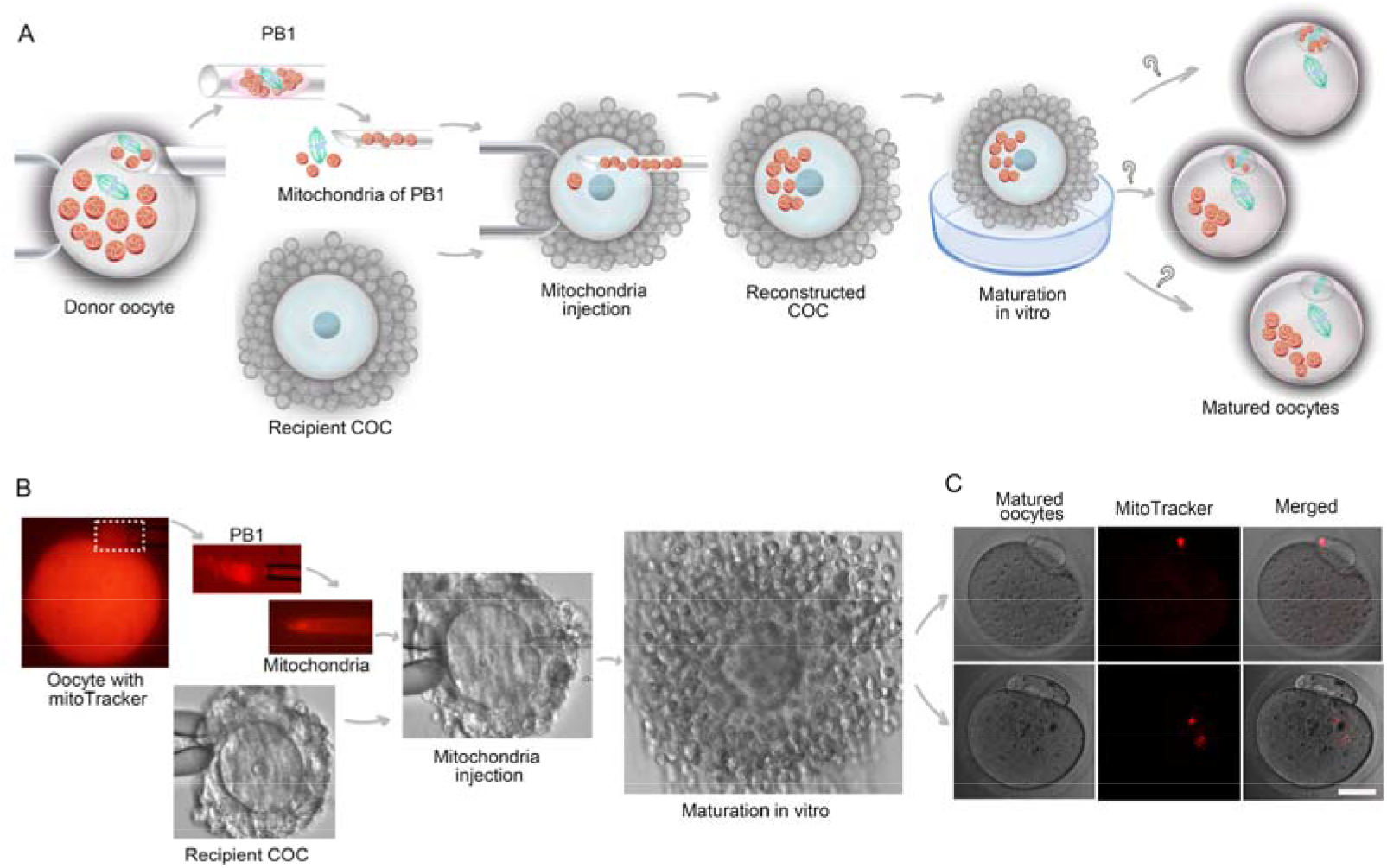
Meiosis I Selected defective Mitochondria into PB1. A. Schematic charts of experimental design of defective mitochondria transfer at GVCOCs and broken GVCOCs stage in this study. B. Micromanipulation of defective mitochondria transfer in GVCOCs in this study. Donor oocyte stained with MitoTracker red, then mitochondria (red) in PB1 was isolated and injected into GVCOCs or broken GVCOCs. Subsequently, the COC was matured in vitro for 10 or 4~5 hours for the GVCOC or broken GVCOCs, respectively. C. Red mitochondria distribution in the PB1 and its oocyte from the matured COCs. Scale bar=40μm.

### The Defective Mitochondria Transfer in Meiosis II Oocyte Suggested Meiosis II Continue to Select Defective Mitochondria into PB2

Unlike PB1, PB2 can survive to the blastocyst stage (Motosugi, 2005), suggesting that mitochondria in PB2 may be consistent with mitochondrial characteristics in the cytoplasm of fertilized egg. Therefore, we did not sequence the mtDNA genome of PB2. And the consistency between the two has also led us to explore whether PB1 is the final destination of defective mtDNA and whether meiosis II will continue to select defective mitochondria to enter PB2.

We first investigated whether the mitochondrial characteristics in PB2 are the same as those in the fertilized eggs of mouse and human. Although TMRE and DilC_1_ (5) were visible in mouse and human PB2 (Figure. S1A,C), Rhod 2 and Fluo-3 were rarely seen in mouse and human PB2s (Figure. S1A,C). The fluorescence intensity showed that there were slight differences for defective mitochondria between mouse PB2 and its sister egg, human PB2 and its sister egg (Figure. S1B,D. Mouse, Dilc5 p=0.0003, TMRE p=0.0011, Rhod2 p=0.0066, Fluo-3: p=0.5196. Human, Dilc5 p=0.0167, TMRE p=0.0362, Rhod2 p=0.0264, Fluo-3: p=0.0603). Compared with the differences between PB1 and its oocyte, these slight differences between PB2 and its sister egg indicated that meiosis II may continue to select defective mitochondria into PB2 if meiosis I did not fully select the defective mitochondria into PB1.

Next, in order to detect the behaviour of the meiosis II for germline selection, defective mitochondria along with the nuclei in mouse PB1 was transferred into an enucleated mouse oocyte to form a reconstructed oocyte, in accordance with PB1 transfer (PB1T) (Wang et al., 2014). Donor mitochondria of mouse oocytes were labelled with 250nM MitoTracker Red. Then PB1T was performed between the stained oocytes and unstained oocytes (Figure. 4A, B, Movie S2). Normally, the nucleus is surrounded by the most active mitochondria due to the nuclear dynamics require energy (Detmer 2007). Thus, to further confirm whether the mitochondria from PB1 are active or defective, we first observed the distribution of mitochondria relative to the nucleus in recombinant oocytes. Eighty-nine recombinant oocytes were obtained from PB1T. Three configurations of donor mitochondria (red mitochondria) distribution related to the nucleus were found in these recombinant oocytes, including front, unilateral, and scattered (Figure. 4C, Table.S4), which strongly suggests that mitochondria in PB1 would be defective. After in vitro fertilization of the recombinant oocytes, a red mitochondrial distribution was observed between the PB2 and its embryos at 2cell stage, as adhered sperm could affect observing fertilized egg at 1-cell stage under the microscope. It is posited that these defective mitochondria have three destinations: completely retained in the embryo, completely extruded along with PB2, or partially retained in the embryo and partially released into PB2 (Figure. 4A). After in-vitro fertilization, Seventy-seven fertilized eggs and Seventy-two embryos of 2-cell were obtained. Confocal imaging showed that red mitochondria were fully extruded into PB2 in three configurations of PB1T recombinant oocytes (Figure. 4D, Table.S3). To rule out the possibility of selecting donor mitochondria and extruding them into foreign organelles, MitoTracker treated ooplast was transferred into oocytes to form a recombinant oocyte. After *in vitro* fertilization, we found that all MitoTracker treated cytoplasm retained in 25 recombinant oocytes (Figure. S2, Table.S3). These results suggest that the meiosis II also has the ability to select defective mitochondria into PB2 if oocyte contained defective mitochondria.

**Figure. 4.**
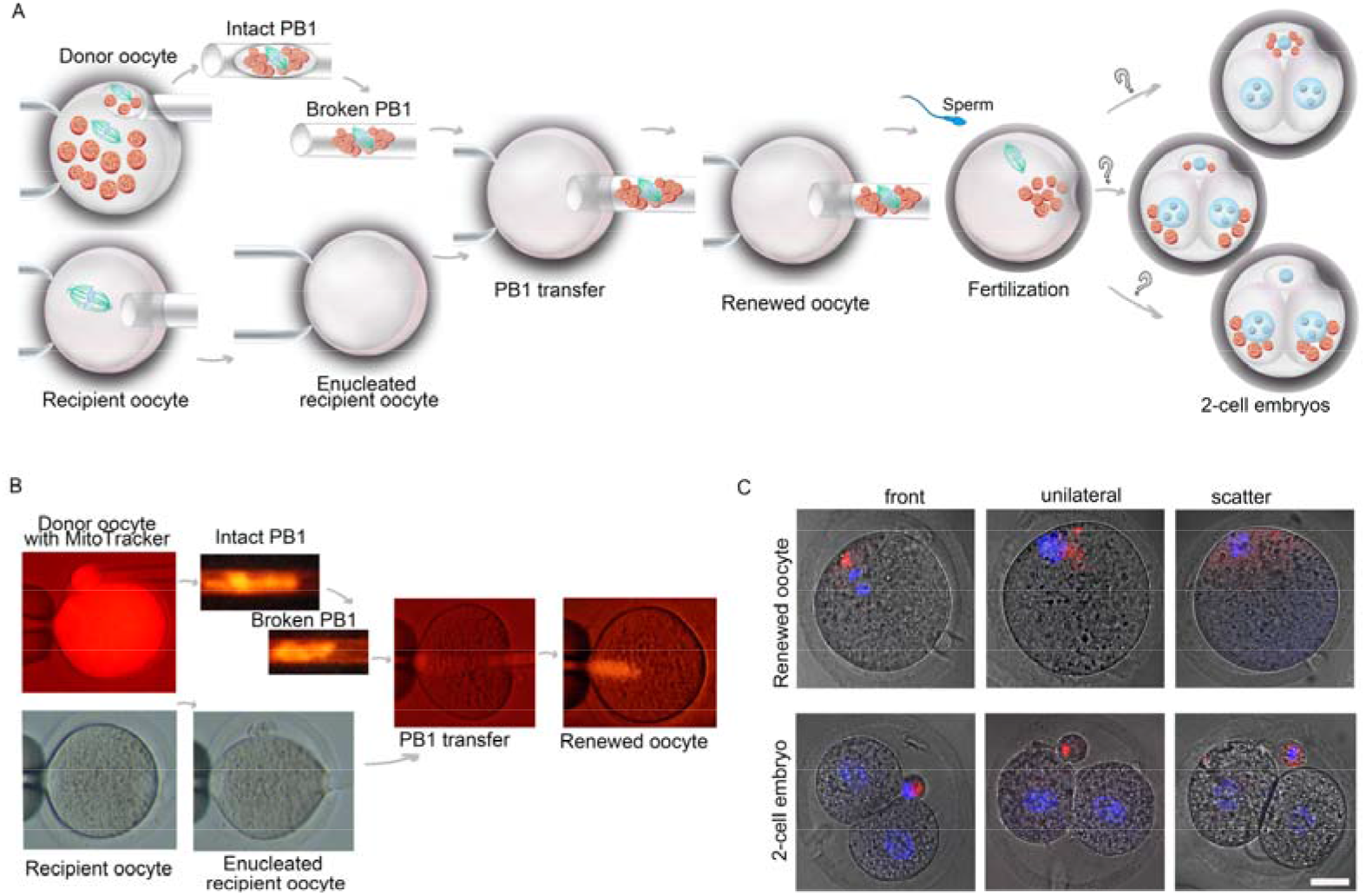
Meiosis II Continue to Select defective Mitochondria into PB2. A. Schematic charts of experimental design of mitochondria transfer at prophase of meiosis II in this study. B. Micromanipulation of mitochondria transfer at prophase of meiosis II in this study: Donor oocyte stained with MitoTracker red, PB1 isolation, recipient oocyte and enucleated oocyte, recombinant oocyte with PB1 genome and mitochondria (red). C. top row: Recombinant oocytes from mitochondria transfer, including three configurations of donor mitochondria (red mitochondria) distribution relative to donor nuclei: front, scattered and unilateral type. Next layer: embryos at 2-cell stage from the recombinant oocytes. Red mitochondria in all type of PB1T recombinant oocytes were totally expelled into PB2. Scale bar=40μm.

## DISCUSSION

There is accumulating evidence to support the occurrence of mtDNA purifying selection during oogenesis (Lieber et al., 2019, Wei et al., 2019, Fan et al., 2008, Stewart et al., 2008, Shoubridge et al., 2008). This process is of great importance in preventing human mitochondrial disease, as the selection can eliminate deleterious mtDNA mutations that escape from genetic bottlenecks, and prevent the offspring from inheriting deleterious mtDNA. Although two studies have provided sequencing evidence in supporting the existence of mtDNA purifying selection during the two meiosis, no intuitive experimental data confirms that meiosis I and II have the function of selecting mutant mtDNA. In addition, the variation of sequencing coverage is too fluctuating among samples owing to the low content of mtDNA in PBs in these studies (De Fanti et al., 2017, Gianoarli et al., 2014). Based on almost 100% mtDNA sequencing coverage, our study demonstrated that mitochondria in PB1 accumulate more mtDNA variants and are defective than mitochondria in their sister oocyte. Furthermore, defective mitochondria transfer in mouse germline showed that both meiosis I and II can extrude defective mitochondria with mutant mtDNA into PB1 and PB2, which strongly enhances the occurrence of mtDNA germline selection at meiosis level.

For the failure to excrete the transplanted mitochondria into PB1 in nine oocytes from GVCOCs group and all oocytes from broken GVCOCs group, we speculated that these GVCOCs were in the advanced GV or MI stage. As we know, cytoplasmic streaming is important for the transport of maternal factors, leading to spindle migration and the establishment of oocyte polarity related to the eventual embryonic anterior-posterior polarity during oogenesis (Duan et al.,2019, Gutzeit et al., 1982, Wolke et al., 2007). We speculated the defective mitochondria was transported with the streaming and the migration of the spindle to the side where the meiosis I would occur in the future. When the meiosis I happened, defective mitochondria with half of the chromosomes extruded to form PB1. When the transplanted mitochondria entered the ooplast of advanced GV or MI oocytes, they might miss the role of cytoplasmic streaming. Then the transplanted mitochondria could not be transported with the migration of the spindle to the side, resulting in the failure to excrete the transplanted mitochondria into PB1.

Hereditary mitochondrial diseases cause a range of serious diseases that can be potentially fatal^1^. It has been noticed that trace amounts of pathogenic mtDNA are conserved in mitochondria replacement, and subsequently outcompete denoted healthy mtDNA, resulting in a reversal to a pathogenic genotype, also known as mitochondrial reversion (Connor 2017, Greenfield 2017). Recently, we have shown that PB1T has the potential to prevent the transmission of mitochondrial diseases from mothers to children and their offspring, as PB1T resulted in undetectable levels of “mutant” mitochondria in mice (Wang et al., 2014, Koch et al., 2014). In this study, our results further provide strong evidence that PB1T has greater potential to thwart mitochondrial reversion in mitochondria replacement, as mitochondria in PB1 is defective and meiosis II can totally squeeze out the mitochondria into PB2 after PB1T following *in vitro* fertilization. Thus, this study also proves the safety and effectiveness of PB1T as it has the potential to prevent mitochondrial reversion in mitochondria replacement.

## AUTHOR CONTRIBUTIONS

H.S. supervised and designed the experiments. H.S. and D.J manipulated mitochondrial transfer. J.Pan. manipulated mtDNA sequence. Y.Yang., S.S and J.Pan. performed staining and confocal analysis. S.S supervised mice and collected oocytes. H.S. and Y.Yang. prepared the figures and wrote the manuscript

## ACKNOWLEDGMENT

This study was supported by grants from National Natural Science Foundation of China (National NSF grants 31871506, Basic key projects of Shanghai Science and Technology Commission 19JC1411200 and 81471512 to H.S.). The authors declare no conflict of interests.

## MATERIALS AND METHODS

### Human oocytes and egg donations

This study was approved by the Institutional Review Board of the First Affiliated Hospital (research license 20160022, 20140222), Anhui Medical University. All the infertile patients agreed and signed informed consent in advance. Totally, 11 patients, who were undergoing regular in-vitro fertilization (IVF) at our reproductive center, donated 25 oocytes for sequencing in this study. Another 15 patients donated 12 oocytes and 12 zygotes for MitoTracker probes staining. All human oocytes in this study were (those) immature oocytes from controlled ovarian hyperstimulation (COH). Since these oocytes must be denuded for ICSI, the meiotic status of these denuded oocytes can be investigated on the morphological grading under microscopic evaluation. All retrieved oocytes were categorized as metaphase II (MII) oocytes, metaphase I (MI), and germinal vesicle (GV). All mature metaphase II (MII) oocytes were used for conventional ICSI. Oocytes at MI stage were donated for this study (Zou et al, 2019). Oocytes at MI stage were maturated in vitro for about 24 hours and monitored under microscopy to observe PB1 extrusion. As soon as PB1 extrusion for about 2 hours, oocyte and its PB1 were isolated for mtDNA sequencing.

### Animals

B6D2F1 (C57/BL6×DBA) was used in this study. All mice used in this study were maintained in accordance with the guidelines of the Laboratory Animal Service, Fudan University (research license 20160225-103).

### Media for manipulation and culture of oocytes and embryos

In this study, the media for the manipulation and culturing oocytes and zygotes were from Vitrolife Sweden AB, Goteborg, Sweden.

### mtDNA genome sequencing

Zone pellucida of oocytes was digested by 0.5% pronase (Roche, 70229227) in 37°C for 5 min. The single oocyte or polar body was sorted into 4 μl PBS. Then 3 μl buffer D2 was added and incubated at 65 °C for 10 min, followed by adding 3 μl stop solution. The lysised products of oocytes were diluted 1:10 and polar bodies were not. Then each PB1 sample was amplified with whole genome amplification (WGA) using REPLI-g Single Cell Kit (QIAGEN, 150345). For each oocyte sample, one-tenth of lysised product was applied for amplification. The DNA concentration was determined by Infinite 200 PRO NanoQuant (TECAN) and diluted to 150ng/μl. Then mtDNA genome of each sample was amplified in 50 μl reaction volume containing 10 μl 5X GC buffer, 1.5 mM Mg^2+^, 200 nM dNTP, 1 μM of each primer, 0.2 μl (1 unit) Taq DNA Polymerase and 1μl template using 9 pair primers (A-H’) (Table S1) (Bannwarth et al., 2009). The PCR was under the following condition: 95°C for 5 min; 38 cycles with denaturation at 95°C for 20 s, annealing at 57°C for 30 s, and elongation at 72°C for 1 min; 1 cycle at 72°C for 10 min with a final 25°C for 10 s (Bannwarth et al., 2009). After estimated by electrophoregram, the PCR products were sequenced using 3730xl DNA Analyzer (ABI). Electropherograms were inspected and aligned to the revised Cambridge reference sequence (NC_012920) using Sequencher 5.4.5. For those mtDNA regions that failed to be amplified or sequenced, PCR reactions were repeated (Figure S1), or new primers (F1-F8, H1-H8) were designed to amplify the failed regions (Table S1). The PCR with F1-F8 and H1-H8 primers under the following condition: 95°C for 5 min; 38 cycles with denaturation at 95°C for 15 s, annealing at 57°C for 15 s, and elongation at 72°C for 30 s; 1 cycle at 72°C for 7 min with a final 25°C for 10s.

For two mix samples of twelve oocytes and twelve PB1s in this study, their lysised products were directly applied to amplify mtDNA genome without WGA

### Sequence Analysis

A Plasmid Editor (APE) software was used to align mtDNA sequences of every two samples. Nucleotides that failed to be sequenced were marked as ‘N’ in sequences. The Mitomap (www.mitomap.org) was used to identify locus of variants and make the functional annotation of AAs change and associated diseases previously reported.

### Membrane potential detection of Mitochondrial

Mouse oocytes were collected at hCG 12.5 to 14 hours. TMRE, DilC 5, Rhod-2 and Fluo Calcium Indicator (Fluo-3 AM) were applied to detect Mitochondrial membrane potential. For TMRE (ab113852, Abcam), DilC_1_(5) (M34151, life Technology) and Rhod-2 (M34151, life Technology) staining, MitoTracker (MitoTracker Green FM, M7514, Life Technology) were used to locate the mitochondria. For Fluo-3AM, MitoTracker (MitoTracker Red CMXRos, M7512, Life Technology) was applied to located the mitochondria. Live oocytes were styed at 37°C for 30 minutes and incubated in G-gamete buffer at 37°C for 20mins afterwards. Images were taken with leica confocal scanning microscope.

### The defective mitochondria transfer in mouse germline Mitochondria transfer at Germinal vesicle stage

Donor mouse oocytes with PB1 were retrieved at HCG 12 hours, cumulus-oocytecumulus complexes were released from ovarian follicles into G-gamete. Cumulus cells were denuded by 3 minutes at incubation with 0.1% hyaluronidase (Sage IVF). Denuded Oocytes with PB1s were stained with 250nM MitoTracker (MitoTracker Red CMXRos, M7512, Life Technology) for 0.5 hour. The mitochondria in PB1 were isolated with 10 and 5.5μm noodle.

Recipient cumulus-oocyte-complexes at GV stage (GVCOCs) were retrieved from ovarian follicles into G-gamete at HCG 5 hours. Then one cumulus-oocyte-complex at GV stage (GVCOCs) or broken GV stage was selected and fixed to the holding needle by applying a vacuum, the mitochondria in 5.5μm noodle were transferred into the ooplast of GVCOCs using microinjection manipulation via 3 clock direction. Subsequently, the GVCOCs and broken GVCOCs were matured in vitro for 10 and 5 hours, respectively. Images of matures oocytes were taken with leica confocal scanning microscope. See also supplemental movie 1.

### Mitochondria transfer at the second meiosis

Cumulus-oocyte-cumulus complexes were released from both ovarian follicles (donor oocytes) and oviducts (recipient oocytes) into G-gamete. Cumulus cells were denuded by 3 minutes at incubation with 0.1% hyaluronidase (Sage IVF). Denuded Oocytes were cultured in G1 medium at 37.5°C, in 5% CO2, 5% O2, and 90% N2 incubation for 30 min before further treatments. Donor mouse oocytes with PB1 were dyed with 250nM MitoTracker. Then mitochondria transfer was performed between PB1 of stained donor oocytes and unstained recipient oocytes using the first polar body genome transfer (PB1T) (Wang et al., 2014). Then PB1T oocytes were fertilized in vitro. Detailed methods for PB1T and IVF were processed according to the previously described methods (Wang et al., 2014). Images of 2-cell embryos were taken with leica confocal scanning microscope. See also supplemental movie 2.

### The active mitochondria transfer (ooplast transfer) in mouse germline

Mouse oocytes at meiosis II were dyed with 250nM MitoTracker. Then ooplast transfer was performed between stained mouse oocytes and unstained mouse oocytes to form reconstructed oocyte. Then reconstructed oocytes were fertilized in vitro. Detailed methods for ooplast transfer and IVF of the reconstructed oocytes were processed according to the previously described methods (Wang et al., 2014). See also supplemental movie 3. Images of 2-cell embryos were taken with leica confocal scanning microscope.

### Statistical Analysis

GraphPad Prism 7 was used to conduct the statistical data analysis. Unpaired t test was performed for the relative fluorescence intensity of the relative fluorescence intensity of human and mouse PB1 to their sister oocyte, human and mouse PB2 to their sister egg, where the significance was set at p < 0.05 (* represents p < 0.05, ** represents p < 0.01, *** represents p <0.001, **** represents p < 0.0001).

